# Theropod dinosaurs had primate-like numbers of telencephalic neurons

**DOI:** 10.1101/2022.06.20.496834

**Authors:** Suzana Herculano-Houzel

## Abstract

Understanding the neuronal composition of the brains of dinosaurs and other fossil amniotes would offer fundamental insight into their behavioral and cognitive capabilities, but brain tissue is only rarely fossilized. However, when the bony brain case is preserved, the volume and therefore mass of the brain can be estimated with computer tomography; and if the scaling relationship between brain mass and numbers of neurons for the clade is known, that relationship can be applied to estimate the neuronal composition of the brain. Using a recently published database of numbers of neurons in the telencephalon of extant sauropsids (birds, squamates and testudines), here I show that the neuronal scaling rules that apply to these animals can be used to infer the numbers of neurons that composed the telencephalon of dinosaur, pterosaur and other fossil sauropsid species. The key to inferring numbers of telencephalic neurons in these species is first using the relationship between their estimated brain and body mass to determine whether bird-like (endothermic) or squamate-like (ectothermic) rules apply to each fossil sauropsid species. This procedure shows that the notion of “mesothermy” in dinosaurs is an artifact due to the mixing of animals with bird-like and squamate-like scaling, and indicates that theropods such as *Tyrannosaurus* and *Allosaurus* were endotherms with baboon- and monkey-like numbers of telencephalic neurons, respectively, which would make these animals not only giant but also long-lived and endowed with flexible cognition, and thus even more magnificent predators than previously thought.

## Introduction

The modern mammal and bird-rich amniote fauna arose from the opportunity created by the demise, in a catastrophic astronomical event, of the giant archosaur species that dominated the Earth during the Mesozoic (Alvarez et al., 1980; Bininda-Emonds et al., 2007; Yu et al., 2021). Until then, sauropodmorphs (the long-necked, plant-eating, quadruped dinosaurs), theropods (the bipedal, carnivorous dinosaurs) and ornithischians (the sometimes armored, plant-eating dinosaurs; Langer et al., 2017) were the largest animals on land. They also had the largest brains amongst land animals at the time, approaching 200 g (the size of a lion’s brain) in sauropodmorphs like *Giraffatitan brancai (Brachiosaurus*) and surpassing that in theropods such as *Tyrannosaurus rex* (up to 343 g; Hurlburt, 1996; Balanoff et al., 2013; Table 1). However, the fact that dinosaurs were sometimes gigantic has been used to imply that their rather large brains were actually undersized for their bodies, giving those animals very low encephalization quotients (EQ) – the first metric to set humans apart from other species, above all others, with a much larger brain than expected for our body size (Jerison, 1973). With low EQs, large dinosaurs could presumably not be that cognitively competent (Jerison, 1973; Hopson, 1977; Knoll and Schwarz-Wings, 2009; Rowe et al., 2011; Ksepka et al., 2020; Balanoff et al., 2013). Rather, it was small theropods such as troodontids, with relatively larger brains and thus higher EQs, that were considered to be “smarter” (Jerison, 1973).

**Table 1.**
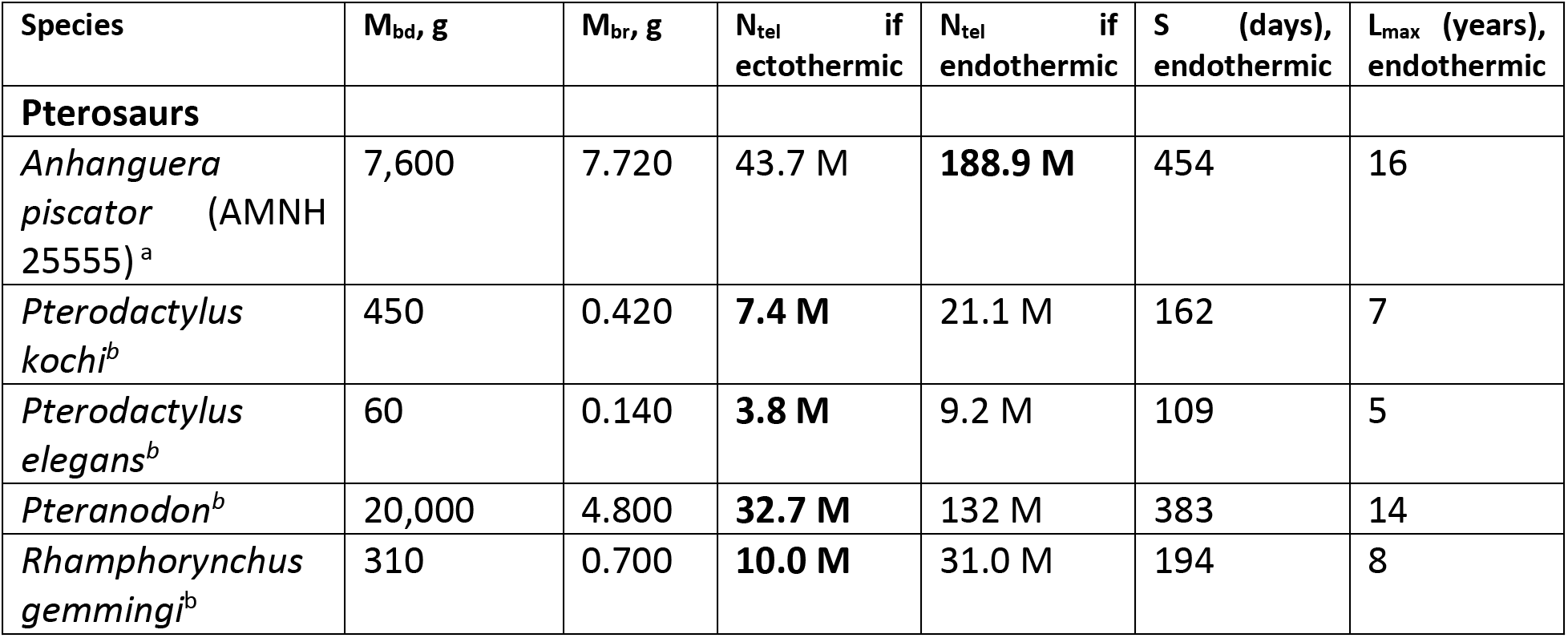
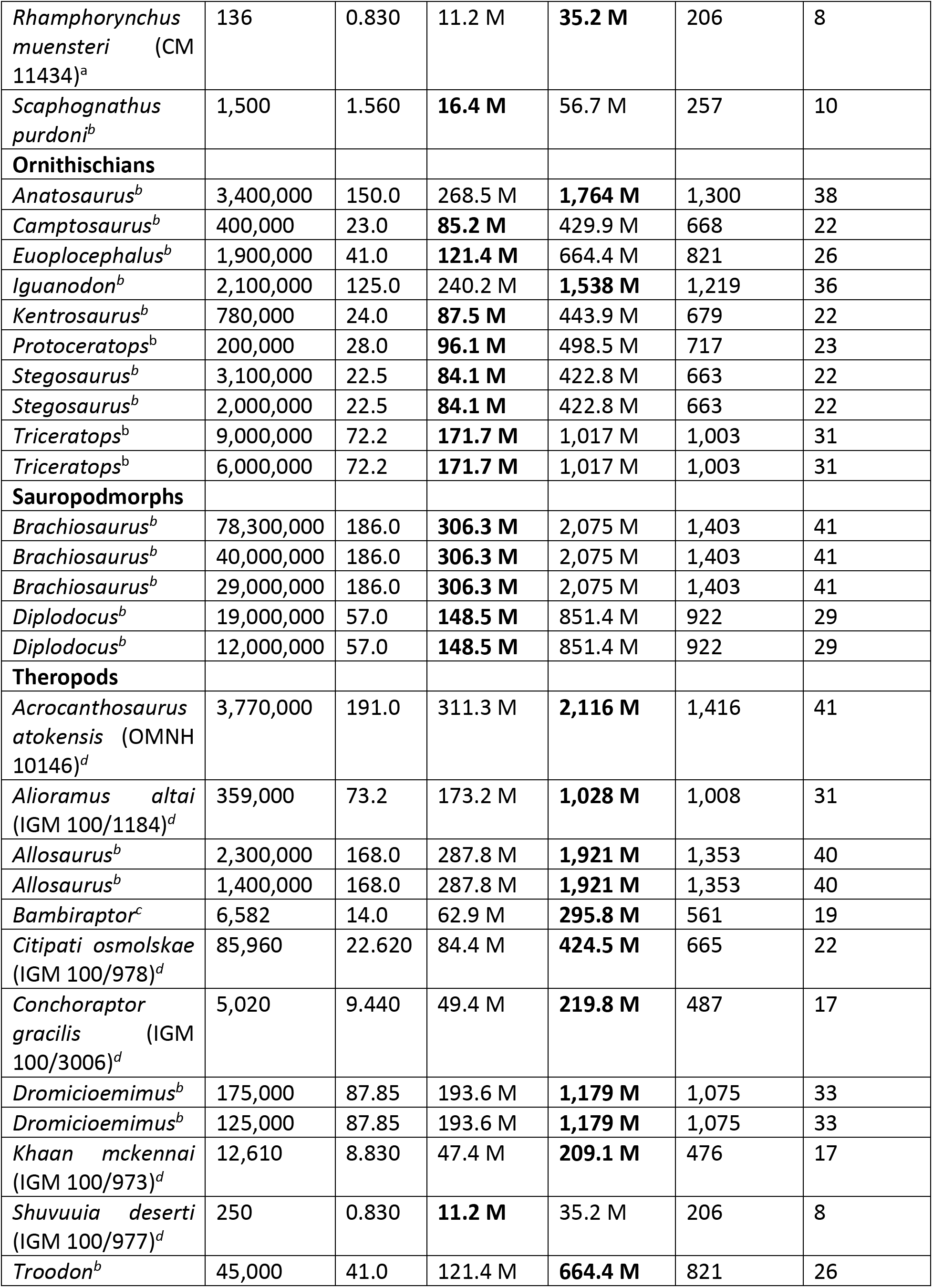
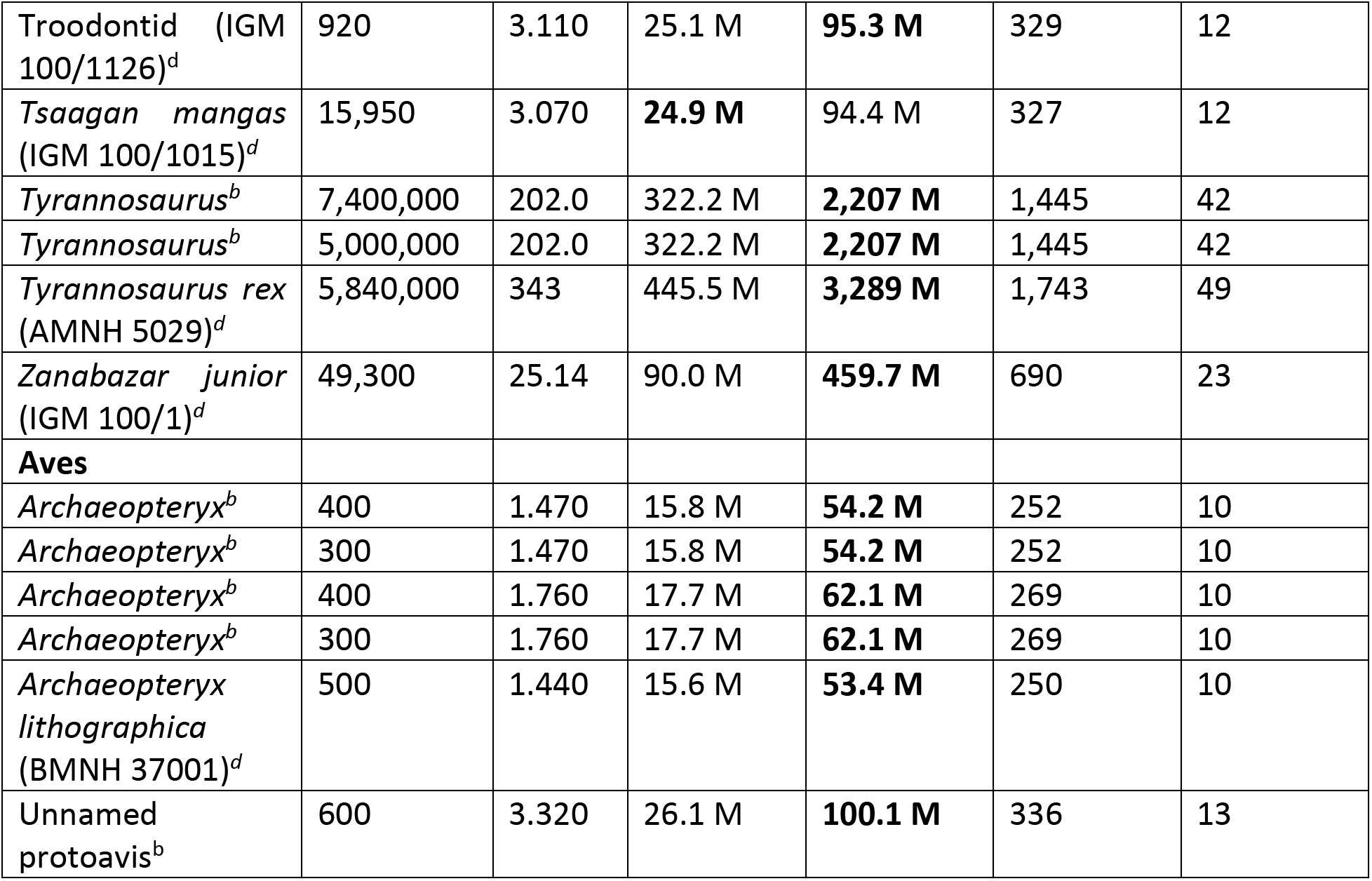
Dataset and numbers of telencephalic neurons (N_tel_) in dinosaur and pterosaur species predicted from brain mass (M_br_) estimates reported in the literature. For the sake of clarity and reproducibility, all data compiled are listed, instead of calculating averages for each species, and are plotted in Figure 2a,b. For each species, estimated N_tel_ (in millions, M) are shown calculated according to the scaling relationships N_tel_ = e^17.518^ M_br_^0.753^ (for endothermic, basal birds) and N_tel_ = e^16.342^ M_br_^0.612^ (for ectothermic reptiles; Figure 1c). Mbd, body mass in grams; M_br_, brain mass in grams, converted from estimated brain volume in cm^3^ using 1 cm^3^ = 1 g. Source of M_bd_ and M_br_ data is indicated next to each species (*a*, Witmer et al., 2003; *b*, Hurlburt, 1996; *c*, Hurlburt et al., 2013; *d*, Balanoff et al., 2013). Values of N_tel_ in bold are the predictions according to Figure 2b.

| Species | $M_{bd}$, g | $M_{br}$, g | $N_{tel}$ if ectothermic | $N_{tel}$ if endothermic | S (days), endothermic | $L_{max}$ (years), endothermic |
| --- | --- | --- | --- | --- | --- | --- |
| **Pterosaurs** |  |  |  |  |  |  |
| *Anhanguera piscator* (AMNH 25555)<sup>a</sup> | 7,600 | 7.720 | 43.7 M | **188.9 M** | 454 | 16 |
| *Pterodactylus kochi*<sup>b</sup> | 450 | 0.420 | **7.4 M** | 21.1 M | 162 | 7 |
| *Pterodactylus elegans*<sup>b</sup> | 60 | 0.140 | **3.8 M** | 9.2 M | 109 | 5 |
| *Pteranodon*<sup>b</sup> | 20,000 | 4.800 | **32.7 M** | 132 M | 383 | 14 |
| *Rhamphorhynchus gemmingi*<sup>b</sup> | 310 | 0.700 | **10.0 M** | 31.0 M | 194 | 8 |
| *Rhamphorynchus muensteri* (CM 11434)<sup>a</sup> | 136 | 0.830 | 11.2 M | **35.2 M** | 206 | 8 |
| *Scaphognathus purdoni*<sup>b</sup> | 1,500 | 1.560 | **16.4 M** | 56.7 M | 257 | 10 |
| **Ornithischians** |  |  |  |  |  |  |
| *Anatosaurus*<sup>b</sup> | 3,400,000 | 150.0 | 268.5 M | **1,764 M** | 1,300 | 38 |
| *Camptosaurus*<sup>b</sup> | 400,000 | 23.0 | **85.2 M** | 429.9 M | 668 | 22 |
| *Euoplocephalus*<sup>b</sup> | 1,900,000 | 41.0 | **121.4 M** | 664.4 M | 821 | 26 |
| *Iguanodon*<sup>b</sup> | 2,100,000 | 125.0 | 240.2 M | **1,538 M** | 1,219 | 36 |
| *Kentrosaurus*<sup>b</sup> | 780,000 | 24.0 | **87.5 M** | 443.9 M | 679 | 22 |
| *Protoceratops*<sup>b</sup> | 200,000 | 28.0 | **96.1 M** | 498.5 M | 717 | 23 |
| *Stegosaurus*<sup>b</sup> | 3,100,000 | 22.5 | **84.1 M** | 422.8 M | 663 | 22 |
| *Stegosaurus*<sup>b</sup> | 2,000,000 | 22.5 | **84.1 M** | 422.8 M | 663 | 22 |
| *Triceratops*<sup>b</sup> | 9,000,000 | 72.2 | **171.7 M** | 1,017 M | 1,003 | 31 |
| *Triceratops*<sup>b</sup> | 6,000,000 | 72.2 | **171.7 M** | 1,017 M | 1,003 | 31 |
| **Sauropodmorphs** |  |  |  |  |  |  |
| *Brachiosaurus*<sup>b</sup> | 78,300,000 | 186.0 | **306.3 M** | 2,075 M | 1,403 | 41 |
| *Brachiosaurus*<sup>b</sup> | 40,000,000 | 186.0 | **306.3 M** | 2,075 M | 1,403 | 41 |
| *Brachiosaurus*<sup>b</sup> | 29,000,000 | 186.0 | **306.3 M** | 2,075 M | 1,403 | 41 |
| *Diplodocus*<sup>b</sup> | 19,000,000 | 57.0 | **148.5 M** | 851.4 M | 922 | 29 |
| *Diplodocus*<sup>b</sup> | 12,000,000 | 57.0 | **148.5 M** | 851.4 M | 922 | 29 |
| **Theropods** |  |  |  |  |  |  |
| *Acrocanthosaurus atokensis* (OMNH 10146)<sup>d</sup> | 3,770,000 | 191.0 | 311.3 M | **2,116 M** | 1,416 | 41 |
| *Alioramus altai* (IGM 100/1184)<sup>d</sup> | 359,000 | 73.2 | 173.2 M | **1,028 M** | 1,008 | 31 |
| *Allosaurus*<sup>b</sup> | 2,300,000 | 168.0 | 287.8 M | **1,921 M** | 1,353 | 40 |
| *Allosaurus*<sup>b</sup> | 1,400,000 | 168.0 | 287.8 M | **1,921 M** | 1,353 | 40 |
| *Bambiraptor*<sup>c</sup> | 6,582 | 14.0 | 62.9 M | **295.8 M** | 561 | 19 |
| *Citipati osmolskae* (IGM 100/978)<sup>d</sup> | 85,960 | 22.620 | 84.4 M | **424.5 M** | 665 | 22 |
| *Conchoraptor gracilis* (IGM 100/3006)<sup>d</sup> | 5,020 | 9.440 | 49.4 M | **219.8 M** | 487 | 17 |
| *Dromicioemimus*<sup>b</sup> | 175,000 | 87.85 | 193.6 M | **1,179 M** | 1,075 | 33 |
| *Dromicioemimus*<sup>b</sup> | 125,000 | 87.85 | 193.6 M | **1,179 M** | 1,075 | 33 |
| *Khaan mckennai* (IGM 100/973)<sup>d</sup> | 12,610 | 8.830 | 47.4 M | **209.1 M** | 476 | 17 |
| *Shuvuuia deserti* (IGM 100/977)<sup>d</sup> | 250 | 0.830 | **11.2 M** | 35.2 M | 206 | 8 |
| *Troodon*<sup>b</sup> | 45,000 | 41.0 | 121.4 M | **664.4 M** | 821 | 26 |
| Troodontid (IGM 100/1126)<sup>d</sup> | 920 | 3.110 | 25.1 M | **95.3 M** | 329 | 12 |
| *Tsaagan mangas* (IGM 100/1015)<sup>d</sup> | 15,950 | 3.070 | **24.9 M** | 94.4 M | 327 | 12 |
| *Tyrannosaurus*<sup>b</sup> | 7,400,000 | 202.0 | 322.2 M | **2,207 M** | 1,445 | 42 |
| *Tyrannosaurus*<sup>b</sup> | 5,000,000 | 202.0 | 322.2 M | **2,207 M** | 1,445 | 42 |
| *Tyrannosaurus rex* (AMNH 5029)<sup>d</sup> | 5,840,000 | 343 | 445.5 M | **3,289 M** | 1,743 | 49 |
| *Zanabazar junior* (IGM 100/1)<sup>d</sup> | 49,300 | 25.14 | 90.0 M | **459.7 M** | 690 | 23 |
| **Aves** |  |  |  |  |  |  |
| *Archaeopteryx*<sup>b</sup> | 400 | 1.470 | 15.8 M | **54.2 M** | 252 | 10 |
| *Archaeopteryx*<sup>b</sup> | 300 | 1.470 | 15.8 M | **54.2 M** | 252 | 10 |
| *Archaeopteryx*<sup>b</sup> | 400 | 1.760 | 17.7 M | **62.1 M** | 269 | 10 |
| *Archaeopteryx*<sup>b</sup> | 300 | 1.760 | 17.7 M | **62.1 M** | 269 | 10 |
| *Archaeopteryx lithographica* (BMNH 37001)<sup>d</sup> | 500 | 1.440 | 15.6 M | **53.4 M** | 250 | 10 |
| Unnamed protoavis<sup>b</sup> | 600 | 3.320 | 26.1 M | **100.1 M** | 336 | 13 |

The EQ is the ratio between the actual brain volume (or mass; these variables are interchangeable, given the specific density of the brain of approximately 1.0 mg/ml) and the brain volume mathematically expected for a species of a given body mass (Jerison, 1973). Calculating the EQ for a species is done under the assumption that brain and body mass are universally correlated across a wide range of species; interpreting the EQ as informative of cognitive capacities, in turn, assumes that brain mass is a universal predictor of numbers of neurons in the brain, and in the pallium (cerebral cortex, in mammals) in particular, the brain structure that most decidedly confers flexibility and complexity to animal behavior (Jerison, 1973). The problem is that while those were reasonable assumptions until the 2000s, we now know that they are both incorrect.

Recent analyses of large datasets show that brain and body size evolve separately in both mammalian (Bertrand et al., 2022) and bird (Ksepka et al., 2020) evolution, making body size an unreliable universal predictor of brain mass. In parallel, a new line of investigation created in my lab based on an original non-stereological method to count brain cells, the isotropic fractionator (Herculano-Houzel and Lent, 2005), allowed the realization that there is no mandatory, universal relationship between body mass and numbers of brain neurons, not even in the brainstem structures that operate the body (Herculano-Houzel et al., 2014, 2015a; Herculano-Houzel, 2017). Additionally, there is no universal relationship between the size of a brain structure and its numbers of neurons; different scaling relationships apply to different clades of mammals and birds (Herculano-Houzel et al., 2014; Herculano-Houzel, 2016), while scaling relationships are mostly shared across reptiles (Kverkova et al., 2022; Herculano-Houzel, 2022). As a result of the lack of a universal correlation between body mass and brain composition, when numbers of neurons (the signal processing units of circuits) should be the limiting factor that determines the computational capacity of a network (Williams and Herrup, 1980), bringing body mass into comparisons of brain size across species of different clades is more than uninformative; it muddles interpretation, by bringing a confounding factor into the mix.

Rather, simple, absolute numbers of neurons in the pallium (organized as a cortex in mammals) are a much better proxy for cognitive abilities than brain or pallial size, whatever the size of the body (Herculano-Houzel, 2017). These are the signal processing units that make behavior flexible and complex and should thus constitute a primary determinant of signal processing capacity (Williams and Herrup, 1980; Herculano-Houzel, 2017; Ströckens et al., 2022). Absolute numbers of pallial neurons are a better predictor of flexible cognitive control than brain mass or EQ across birds and primates alike (Herculano-Houzel, 2017), and, across bird species, they are a great predictor of innovation rate (Sol et al., 2022). It thus follows that understanding the capability for behavioral and cognitive flexibility of the extinct species that once dominated the Earth’s fauna requires going beyond the veil of brain and body size and gaining direct understanding of the numbers of neurons that composed the pallium of those animals. That understanding can be reached once the scaling relationship between brain size and numbers of neurons for species in a given clade is known, which makes brain size a reliable proxy of numbers of neurons in that clade. In particular, numbers of pallial, or telencephalic, neurons in extant species can be estimated from brain size by clade-specific allometric power functions with r^2^ values of typically 0.9, that is, which provide estimates with about 90% reliability (Herculano-Houzel, 2019a). Using this expedient, we have previously been able to infer the numbers of neurons that composed the brain of prehistoric hominin species (Herculano-Houzel and Kaas, 2011) and fossil mammals (Herculano-Houzel et al., 2011).

How to determine the scaling relationships that applied to fossil species, when brain tissue is not preserved in the fossil record? Brain mass of dinosaur species can be estimated with CT or micro-CT of extant or fossilized skulls (Hulburt, 1996; Knoll et al., 1999; Witmer et al., 2003; Balanoff et al., 2013). In these cases, the numbers of neurons that composed their brains, and their telencephalon in particular, can be estimated if one can determine the applicable predictive equations relating numbers of neurons to brain mass. Dinosaurs and pterosaurs (together with turtles and crocodilians) were archosaurs, the sister clade to modern squamates, which are ectothermic sauropsids; but living birds, which are endothermic sauropsids, are surviving dinosaurs. Thus, the alternative hypotheses tested here are that pterosaur and dinosaur brains were either composed like ancestral, and modern, ectothermic sauropsid brains, or had already shifted to the composition of modern, endothermic, early-branching bird brains (Herculano-Houzel, 2022).

Here I determine whether ectothermic or endothermic sauropsid scaling rules most likely applied to different dinosaur (and pterosaur) species, and use the allometric scaling equations that apply to the brain structures of modern birds and non-avian sauropsids to predict the numbers of neurons that composed the brains of fossil dinosaurs and pterosaurs based on their brain volumes. These allometric scaling equations describe the relationship between brain structure mass and numbers of neurons, determined using the isotropic fractionator (Herculano-Houzel and Lent, 2005). This method, which consists of turning brains or brain structures into a homogeneous soup of floating cell nuclei that allows the fast, unbiased, and reproducible estimation of how many neuronal and non-neuronal cells composed those structures (Herculano-Houzel et al., 2015b), has by now been applied to over 200 species of mammals, birds, and non-avian sauropsids (Kverkova et al., 2022). While the resulting dataset did not separate the pallial from the subpallial structures that together compose the telencephalon of non-avian sauropsids, it did establish that the vast majority of telencephalic neurons are found in the pallium of early-branching birds; thus, numbers of telencephalic neurons in the dataset offer a good approximation for the number of pallial neurons (though the subpallium also contributes to flexible behavior; Boot et al., 2017). In contrast to the recent initial analysis of the full dataset which focused on relationships between numbers of neurons and body mass (Kverkova et al., 2022), here I concentrate on the clade-specific relationships between telencephalic and brain mass and numbers of telencephalic neurons in the different clades of living avian- and non-avian sauropsids in search of establishing what relationships putatively applied to fossil species, which I then use to estimate numbers of telencephalic neurons in select species with known brain volume and mass.

## Methods

All data on numbers of telencephalic neurons, brain and telencephalic mass, and body mass for 174 extant sauropsid species (avian and non-avian) used to calculate the scaling relationships in Figure 1 were taken from Kverkova et al. (2022). All power functions were calculated using least-squares regression of log-transformed values using the JMP 16 software package (Carey, NC). Power functions were calculated separately for different groups of birds, as detailed in the results (Accipitriformes, n=4 species; Anseriformes, n= 7; Columbiformes, n=5; Falconiformes, n=3; Galliformes, n=9; Palaeognathae, n=6 species; Passeriformes, n=13; Psittaciformes, n=11; Strigiformes, n=7); and for all non-avian sauropsids in the dataset (88 squamates, 19 testudines and 1 crocodilian species; Kverkova et al., 2022). Particular emphasis was placed on the early-branching (“pre-K-Pg”) bird species, which in this dataset belong to Palaeognathae, Galliformes, Anseriformes and Collumbiformes, which share scaling relationships (see Figure 1, and Herculano-Houzel, 2022). Thus, all scaling relationships reported pertain to a group of species that belong to closely related clades that share a scaling relationship amongst themselves. *Within* each group, I chose to not adjust the scaling exponent for phylogenetic relatedness in the dataset because the point of calculating these scaling relationships was not to study phylogeny, but to use the mathematical functions to make predictions from brain mass, in which case the desired scaling relationship is the one that applies to the raw data, with no modifications introduced that would distort the mathematical reality of the relationships between the physical entities (structure mass) and (number of neurons).

**Figure 1.**
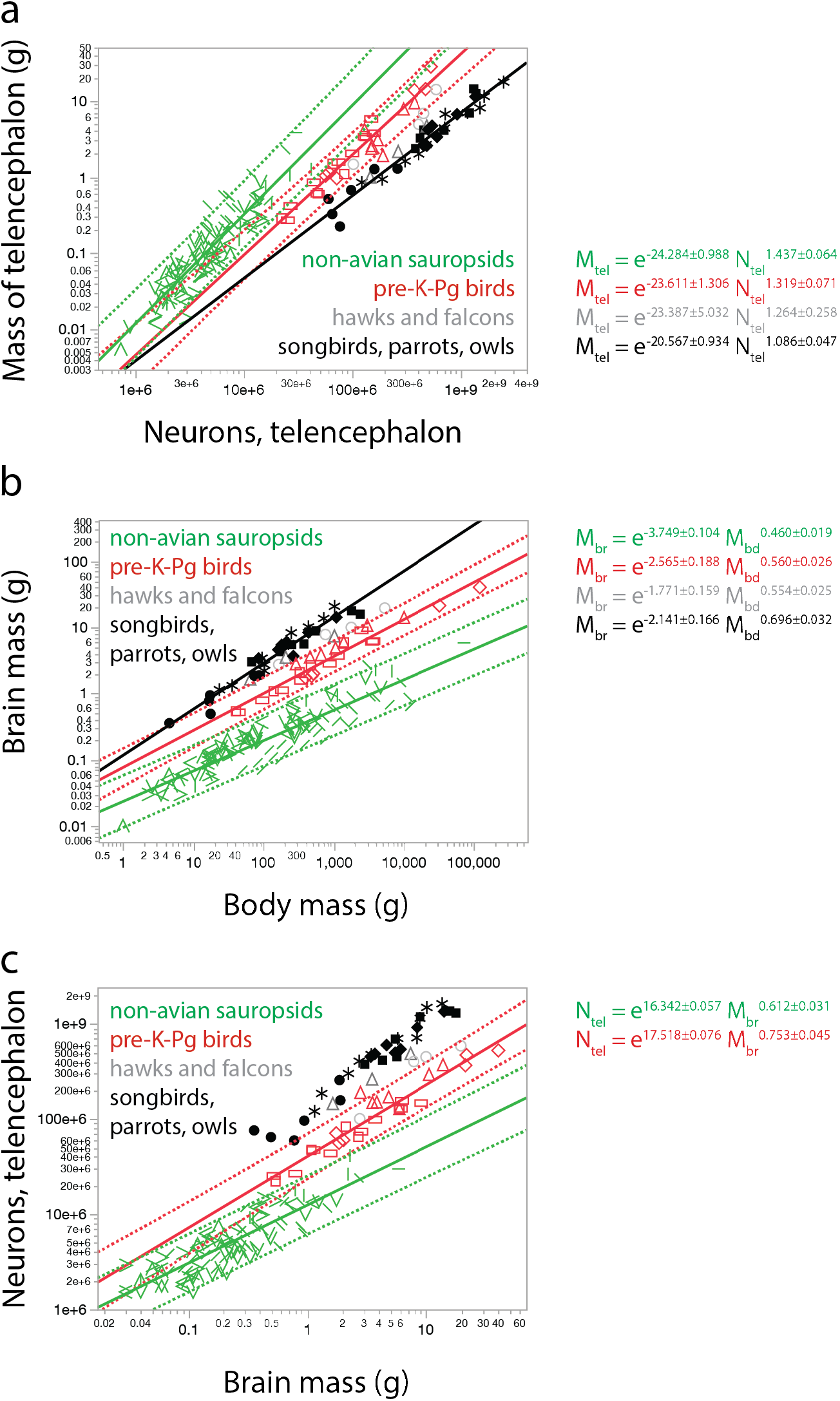
Scaling relationships that apply to extant avian and reptile species. **A**, clade-specific scaling of telencephalic mass (M_tel_) with numbers of telencephalic neurons (N_tel_) distinguishes pre-K-Pg bird clades (Palaeognathae, red lozangles; Galliforms, red rectangles; Anseriformes, red triangles; Columbiformes, red squares) from post-K-Pg bird clades (Falconiformes, grey triangles; and Accipitriformes, grey circles, i.e. hawks and falcons; and Passeriformes, black circles; Psittaciformes, black asterisks; and Strigiformes, black squares, i.e. songbirds, parrots, and owls; and non-avian sauropsids (including Testudines and Crocodilia). Power functions are indicated next to the graph for each group (non-avian sauropsids in green, with different symbols for the different clades in Kverková et al. (2022), r^2^=0.826, p<0.0001, n=108; pre-K-Pg birds in red, r^2^=0.933, p<0.0001, n=27 species; Accipitriformes and Falconiformes, fit not plotted for clarity, r^2^=0.828, p=0.0045, n=7 species; Passeriformes, Psittaciformes and Strigiformes in black, r^2^=0.949, p<0.0001, n=31 species). **B**, clade-specific scaling of brain mass with body mass similarly distinguishes pre-K-Pg bird clades (Palaeognathae, Galliforms, Columbiformes, Anseriformes) from other, later-branching bird clades (Falconiformes and Accipitriformes; and Passeriformes, Psittaciformes, and Strigiformes), and non-avian sauropsids. Power functions as in A: non-avian sauropsids plotted in green, r^2^=0.842, p<0.0001, n=108 species; pre-K-Pg birds plotted in red, r^2^=0.948, p<0.0001, n=27 species; Accipitriformes and Falconiformes, in grey, fit not plotted for clarity, r^2^=0.990, p<0.0001, n=7 species; and Passeriformes, Psittaciformes and Strigiformes plotted in black, r^2^=0.944, p<0.0001, n=31 species. **C**, clade-specific predictive relationships for estimating numbers of telencephalic neurons from brain mass for pre-K-Pg birds and non-avian sauropsids. Power functions are indicated next to the graph for pre-K-Pg birds (in red; r^2^=0.918, p<0.0001, n=27 species) and for non-avian sauropsids (in green: r^2^=0.791, p<0.0001, n=108 species). All data from Kverkova et al., 2022.

Absolute estimated brain mass (not simply endocranial volume) and body mass values in fossil pterosaur and dinosaur species shown in Figure 2 were collected from four studies that compiled estimates from CT studies of endocranial volume (Table 10 in Hulburt, 1996; Witmer et al., 2003; Hulburt et al., 2013; Balanoff et al., 2013). Different brain-to-endocast fractions by definition will impact the estimated brain mass or volume (Knoll et al., 2022); thus, care was taken to use explicitly reported estimates for brain mass, whether or not that matched the endocranial volume. Estimates for theropod and pterosaur species consider that theropod and pterosaur brains filled the endocranial cavity (Balanoff et al., 2013; Witmer et al., 2003), whereas Hurlburt’s estimates for all dinosaurs except small theropods considered that brain mass was half that of endocast volume (Hurlburt, 1996, p. 147). Where available, specimen numbers are provided in Table 1 (they were not reported in Hurlburt’s 1996 dataset). Where more than one estimate was available for a species, all values are plotted so as to allow the evaluation of the impact of specimen and methodological variability, as well as the range of body and brain size combinations. I note that the taxonomy of several fossil species has been updated since Hurlburt’s data compilations that I use in Table 1. For the sake of making the present dataset transparent and readily verifiable by other researchers and useful to them, and because I am not a paleontologist but a neuroscientist using paleontological data, I chose to preserve in Table 1 the original names under which the data were originally reported by the authors.

**Figure 2.**
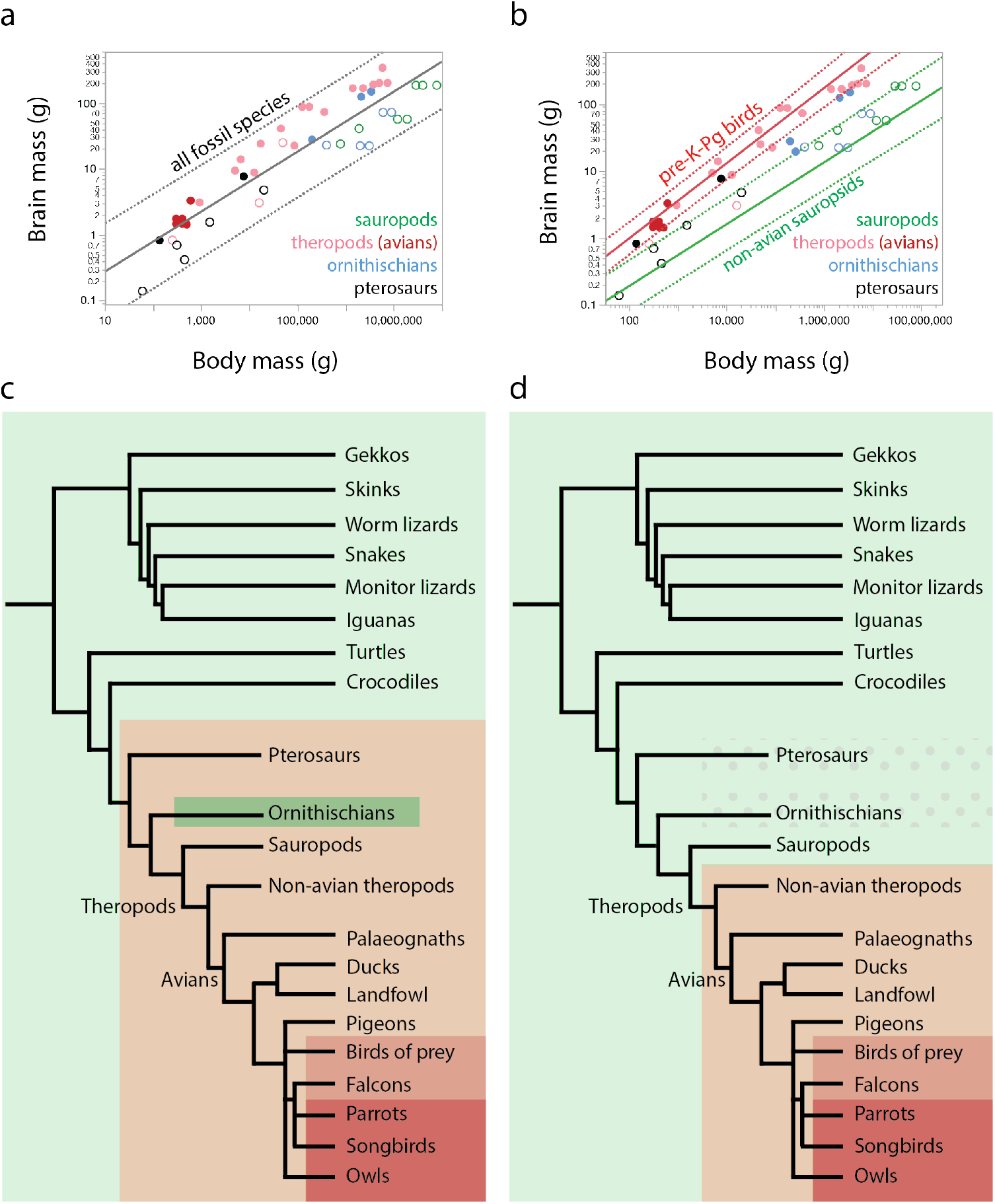
Dinosaur and pterosaur species vary in conforming to either modern pre-K-Pg endothermic (bird) or ectothermic (non-avian sauropsid) scaling relationships between brain and body mass. **A,** A single power function M_br_ = e^-2.327±0.338^ M_bd_^0.455±0.028^ (r^2^=0.857, p<0.0001, plotted in grey) can be fit to the relationship between brain and body mass (M_br_ and M_bd_, respectively) across all the fossil dinosaur and pterosaur species in the dataset (Table 1), color-coded as indicated in the key. However, most non-avian theropod species (pink) have larger brain mass than predicted for their body mass, whereas sauropodmorphs (green) and most pterosaur species (black) have smaller brain mass than predicted by this joint scaling function, which is consistent with a better account of the distribution by two separate functions. Filled circles correspond to species that match brain x body mass scaling in pre-K-Pg birds; unfilled symbols denote species that match brain x body mass scaling in non-avian sauropsids, as in B. **B,** Data points for fossil species in Table 1 plotted onto the fitted power functions shown in Figure 1b that describe the brain x body mass relationship in modern ectothermic sauropsids and pre-K-Pg endothermic sauropsids show that most non-avian theropods (pink) and early avians in the dataset have the brain mass expected for a generic endothermic, pre-K-Pg modern bird that had their body mass, whereas most sauropodmorphs and pterosaurs have brain mass within the range expected for a modern generic ectothermic, non-avian sauropsid of their brain mass. Different ornithischian species conform to one of the other scaling relationship. Power functions, plotted with 95% prediction intervals, are the same as in Figure 1b. **C, D,** schematics of alternate proposals for the evolution of brain vs body mass relationships that are characteristic of ectothermic (green) or endothermic (shades of red) modern amniotes. Evolutionary trees based on Wiemann et al. (2022) and Kverkova et al. (2022). **C,** metabolite-based analysis (Wiemann et al., 2022) predicts that a brain x body scaling relationship similar to that characteristic of modern endothermic pre-K-Pg birds applied broadly across dinosaur and pterosaur species, but not in ornithischians. **D**, present brain x body scaling relationships shown in **B** predicts that endothermy was widespread in theropods but only occasional in pterosaurs and ornithischians.

## Results

Figure 1a shows that the neuronal scaling rules that apply to the telencephalon of modern birds and non-avian sauropsids are clearly distinct. For a similar telencephalic mass (which occurs in the largest non-avian sauropsids and the smallest birds), early-branching bird clades in the dataset (Palaeognathae, Galliformes, Anseriformes, and also Columbiformes, in red), which arose before the K-Pg boundary (henceforth, pre-K-Pg birds; Brusatte et al., 2015), have ca. 5 times as many telencephalic neurons as non-avian sauropsids (in green), and the post-K-Pg-branching Passeriformes, Psittaciformes and Strigiformes clades have even more telencephalic neurons (in black). For instance, the zebra finch has 55 million telencephalic neurons whereas the Sudan plated lizard has only 14 million, though both have a telencephalon of ca. 0.3 g. Strikingly, there is very little overlap in numbers of telencephalic neurons between birds and non-avian sauropsids, a distinction that I have hypothesized to result from the increased oxidative rates that make endothermy possible in birds compared to other sauropsids (Bennett and Ruben, 1979) rather than from endothermy itself (Kverkova et al., 2022; Herculano-Houzel et al., 2022). Figure 1b shows that endothermic sauropsids (i.e., birds) do have larger brains compared to ectothermic sauropsids of a similar body mass, but again in a clade-specific manner, such that bird species belonging to post-K-Pg clades (songbirds, parrots, and owls) have even larger brain mass for a similar body mass.

Together, the distinction between bird clades in Figures 1a and 1b shows that a shift to endothermy cannot be the sole cause of increased brain mass and numbers of telencephalic neurons in birds relative to body mass (Herculano-Houzel, 2022; Gillooly and McCoy, 2014), since brain mass increases further relative to body mass in the post-K-Pg songbirds, parrots and owls compared to the pre-K-Pg birds. Importantly, these findings establish that comparisons across species and clades for where brain scaling is involved cannot treat “birds” as a single entity, as has been standard in the field (Balanoff et al., 2013). However, amongst non-avian sauropsid clades, neuronal scaling rules are much more uniform (Herculano-Houzel, 2022), and for the purposes of this study, all extant non-avian sauropsids in the dataset (88 squamates, 19 testudines and 1 crocodilian species) can be considered to share the scaling rules of interest, which are clearly distinct from the scaling rules that apply to extant pre-K-Pg birds (Figure 1a).

Expressing the number of telencephalic neurons in the brain as a function of brain mass shows that within a clade, brain mass has strongly predictive power to arrive at estimates of numbers of telencephalic neurons in a brain of known mass, once the neuronal scaling rules that presumably apply are known. Figure 1c shows that clearly different predictive scaling rules apply to extant pre-K-Pg birds and to non-avian sauropsids, with non-overlapping 95% prediction intervals across the entire range of bird-like brain sizes. Specifically, these distinct power laws are such that over 80% of the variation in numbers of telencephalic neurons in non-avian sauropsid species, and over 90% in pre-K-Pg birds, can be accounted for by the variation in brain mass, if clade identity is respected.

The predictive equations shown in Figure 1c can be used to infer the numbers of telencephalic neurons that composed the brains of dinosaur, pterosaur, and other fossil sauropsid species provided that these species are found to conform to the scaling rules that apply to either modern pre-K-Pg birds or non-avian sauropsids. Figure 2, using data compiled from the literature (Table 1), shows that the scaling of brain mass with body mass can indeed provide a distinguishing criterion across dinosaur species. Standard practice in the field has been to assume that a single scaling relationship applies homogeneously across mixed dinosaur clades (Jerison, 1973; Balanoff et al., 2013; Grady et al., 2014). Figure 2a confirms that a highly significant single scaling relationship can indeed be fit to the ensemble of the fossil species sampled, with a 95% prediction interval that includes all but one species (Figure 2a), with an exponent of 0.455±0.028 (r^2^=0.857, p<0.0001) that is similar to the exponent that applies to extant non-avian sauropsids only (0.460±0.019, r^2^=0.842 in Figure 1b) but with a significantly larger intercept of e^-2.327±0.338^ compared to the e^-3.749±0.104^ of modern such species. If this joint scaling relationship intermediate between living non-avian sauropsids and birds truly applied across all dinosaurs, as calculated previously (Balanoff et al., 2013), and these diverse species shared a single scaling relationship between brain and body mass (for example, if they were “mesothermic”, as once suggested by a similar expedient of analyzing dinosaur species together regardless of clade; Grady et al., 2014), then it would not be justified to apply the neuronal scaling rules of either modern birds or no-navian sauropsids to these fossil species.

In contrast, Figure 2b shows that individual dinosaur and pterosaur specimens clearly conform to the brain x body scaling rules that apply to either ectothermic or pre-K-Pg endothermic extant sauropsids. Both *Archaeopteryx*, the earliest avian species of known brain and body mass, and a non-identified “protoavis” (Hulburt, 1996; filled red circles) conform to the scaling relationship that applies to modern pre-K-Pg birds, which originated within Jurassic theropods (Brusatte et al., 2015), with brains significantly larger than expected for a modern reptile of similar body mass. Likewise, the majority of theropod dinosaur species of known brain and body mass (filled pink circles) conform to the brain vs body mass relationship that applies to modern pre-K-Pg birds, with the exception of *Shuvuuia deserti* (with brain mass just below the prediction interval for pre-K-Pg birds) and *Tsaagan mangas* (with the brain mass expected for a modern ectothermic sauropsid of similar body mass; unfilled pink circles). Conversely, most sauropodmorph dinosaurs in the dataset had the brain mass expected for a modern ectothermic sauropsid of their body mass (unfilled green circles). Ornithischian (blue circles) and pterosaur species (black circles), in turn, align either with endothermic pre-K-Pg birds (filled circles) or ectothermic sauropsids (unfilled circles) in their brain vs body mass relationship, depending on the species (Figure 2b, Table 1). *Protoceratops* (filled blue circle), in particular, approached the distribution of modern pre-K-Pg birds. Thus, the comparison of the brain vs body mass relationships of the sampled fossil species with those of modern ectothermic and endothermic pre-K-Pg sauropsids suggests that the neuronal scaling rules shared by modern ectothermic sauropsid species also applied to the telencephalon of all non-theropod fossil species of sauropsids, with the exception of some pterosaur and ornithischian species (filled data points in Figure 2b), whereas theropods as a whole already had neuronal scaling rules similar to those of modern, endothermic pre-K-Pg sauropsids, according to the cladogram in Figure 2d. Figure 3a shows how the brain x body mass relationship of fossil avian and non-avian theropods matches that of extant pre-K-Pg bird species (and, in comparison, primates have much larger brains for a similar body mass). Figure 3a also shows that despite the extreme difference in body size, the brain x body mass relationship of fossil sauropodmorphs, several pterosaurs and some ornithischians match the relationship that applies to extant non-avian sauropsids.

**Figure 3.**
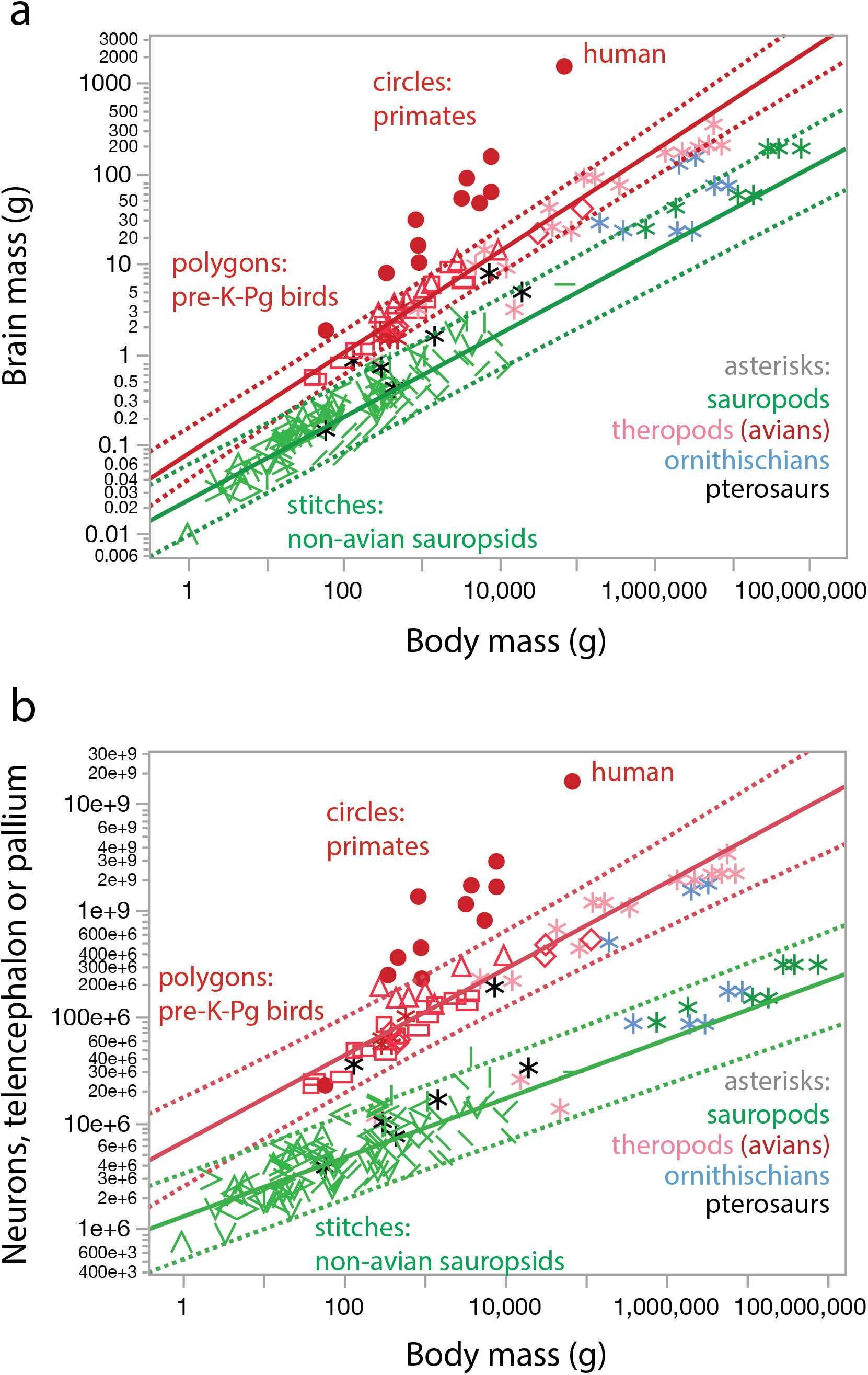
Theropod dinosaurs overlap with modern pre-K-Pg endothermic sauropsids (birds) in their brain x body mass scaling, which results in primate-like numbers of telencephalic neurons in theropods, whereas most sauropodmorphs, pterosaurs and some ornithischians overlap with ectothermic non-avian sauropsids. **A,** Data points for fossil species in Table 1 (asterisks) as well as data points for extant pre-K-Pg bird species (red polygons), extant non-avian sauropsids (green stitches) and primates (red circles) plotted onto the fitted power functions shown in Figure 1b that describe the brain x body mass relationship in modern ectothermic sauropsids and pre-K-Pg endothermic sauropsids. Power functions, plotted with 95% prediction intervals, are the same as in Figure 1b. **B**, Same as in **A**, but showing measured numbers of neurons in the mammalian cortex or in the telencephalon of extant sauropsids plotted together with estimated numbers of neurons in the telencephalon of dinosaur and pterosaur species (bold values in Table 1). Notice the similar numbers of neurons in the telencephalon of theropods and in the cerebral cortex of primates despite much larger body masses in theropods.

Given the striking distinction in brain x body scaling between extant non-avian sauropsids and pre-K-Pg birds shown in Figure 1, most likely associated with the distinction between ectothermy and endothermy (Kverkova et al., 2022; Herculano-Houzel, 2022), and the similar segregation of fossil dinosaur and pterosaur species shown in Figure 2, here I take the approach of hypothesizing that the neuronal scaling rules calculated for the telencephalon of endothermic or ectothermic modern species already applied to the brains of fossil species of matching brain vs body scaling relationship. Thus, considering that most fossil theropods had brains of the mass expected for a modern pre-K-Pg bird of theropod-like body mass (Figure 2b), the predictive neuronal scaling rule calculated for modern pre-K-Pg birds will also estimate the numbers of telencephalic neurons in fossil theropod species of known brain mass.

Using the published values of brain mass estimated from CT analysis (Table 1) plotted in Figure 2, I find that theropods had primate-like numbers of telencephalic neurons (Figure 3B; Table 1), although in bodies as much as 1,000 times larger than primates of similar numbers of cortical neurons. As depicted in Figure 4, numbers of telencephalic neurons ranged from just over 1 billion telencephalic neurons in the 73 g brain of *Alioramus*, comparable to a capuchin monkey, to over 3 billion telencephalic neurons in a 343 g brain of *Tyrannosaurus rex*, which is more telencephalic neurons than found in a baboon (Herculano-Houzel et al., 2015a). In comparison, scaling with ectothermic sauropsid rules, *Triceratops*, with a 72 g brain similar in size to *Alioramus*, presumably had only around 172 million telencephalic neurons – fewer than the 306 million neurons found in the cerebral cortex of a capybara (Herculano-Houzel et al., 2015a). Importantly, the use of endotherm (avian sauropsid) scaling rules to estimate numbers of telencephalic neurons in theropods versus ectotherm (non-avian sauropsid) scaling rules in ornithischians is supported by recent metabolite findings in these species (Wiemann et al., 2022). The distinction is highly consequential: if the *Tyrannosaurus* brain scaled like a non-avian ectothermic sauropsid brain, it would have an estimated 446 million telencephalic neurons – still as many as in a large dog, but less than 15% of the baboon-like 3.3 billion telencephalic neurons estimated if pre-K-Pg bird-like scaling rules applied (Table 1).

**Figure 4.**
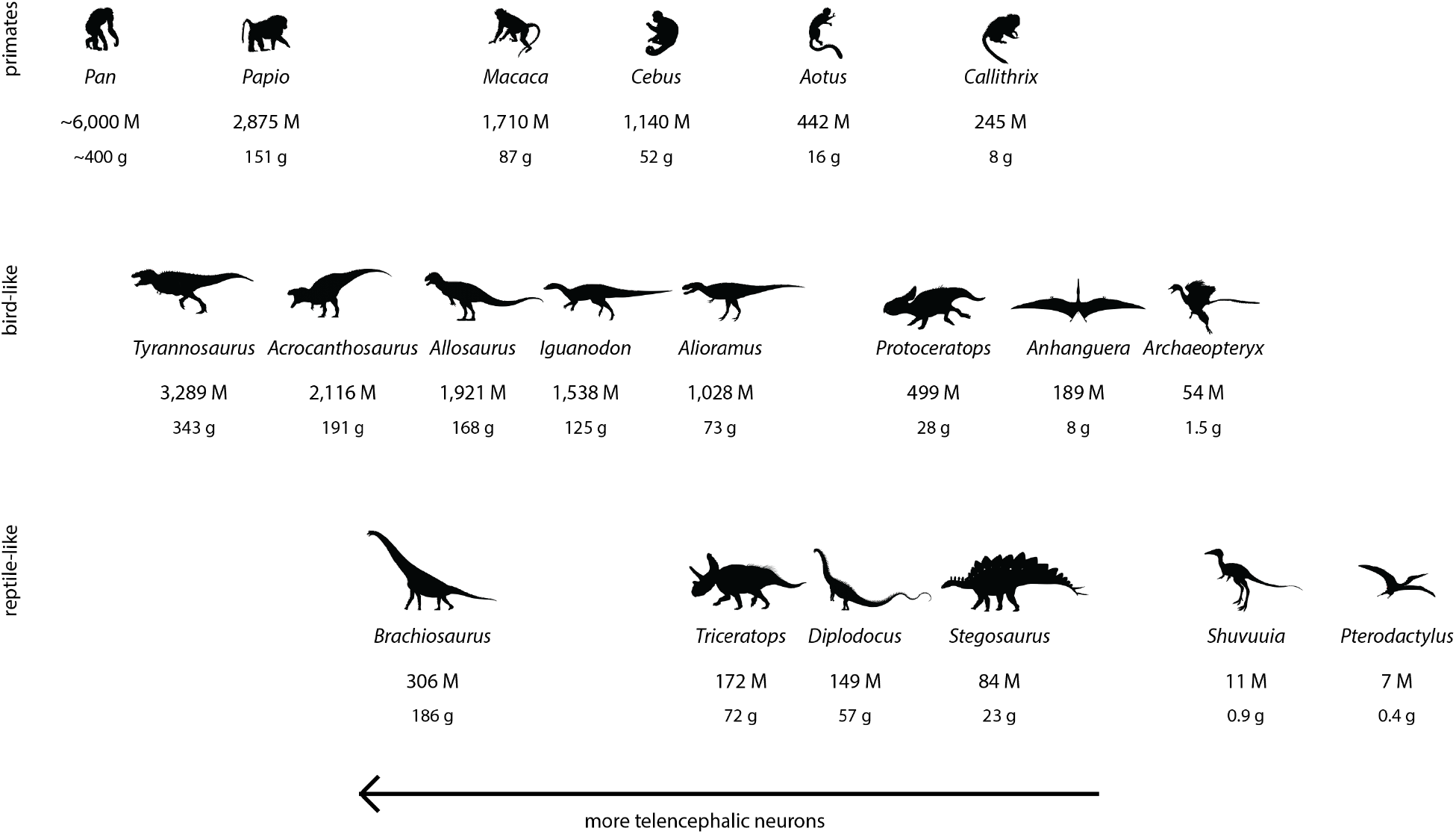
Estimates of numbers of telencephalic neurons in dinosaur and pterosaur species. Values below each image are numbers of telencephalic neurons (in millions, M) and brain mass (in grams). Select species from the dataset in Table 1 are depicted, ranked from left to right in decreasing order of numbers of telencephalic neurons and separated according to whether the species conforms to the brain vs body mass relationship of modern (endothermic) pre-K-Pg birds (center row) or of modern non-avian (ectothermic) sauropsid species (bottom row). For comparison, select primate species of similar numbers of pallial neurons are shown in the top row. All images from phylopic.org.

In pterosaurs, the brain x body relationship in Figure 2b supports the metabolic finding consistent with endothermy in *Rhamphorhynchus muensteri* (though not in *R. gemmingi;* Table 1), but is consistent with ectothermy in *Pteranodon* (Wiemann et al., 2022). Assuming pre-K-Pg endothermic sauropsid scaling rules, the 8 kg pterosaur *Anhanguera* had an estimated 189 million telencephalic neurons, fewer than in a marmoset, in a brain of 8 g, which is almost 4 times as many telencephalic neurons as estimated in the small brain of *Archaeopteryx* (Figure 3). In contrast, assuming ectothermic sauropsid scaling rules, a 450 g *Pterodactylus* animal had only an estimated 7 million telencephalic neurons, fewer than found in a mouse, in its 0.4 g brain (Table 1).

Comparing the present dataset with the species analyzed in that study of fossil metabolites (Wiemann et al., 2022), the only disagreement regards *Diplodocus*, the only sauropodmorph tested in that study, suggested to have elevated oxidative metabolic rates consistent with endothermy but found here to conform to the brain x body scaling relationship of modern ectothermic sauropsids. Employing the latter, according to Figure 2b, sauropodmorphs such as *Brachiosaurus* and *Diplodocus* had only marmoset-like numbers of telencephalic neurons in their brains, even though these had mass that was similar in range to the brains of theropods that had estimated monkey-like numbers of telencephalic neurons (Figure 4, bottom row). Thus, until the metabolite concentrations of sauropodmorph species can be systematically analyzed in more species, the present data suggest that these large quadrupeds had telencephalons that were composed according to the neuronal scaling rules that still apply to modern ectothermic sauropsids - which, incidentally, are the same that apply to modern nonprimate mammals (Herculano-Houzel, 2022).

## Discussion

Here I show that the scaling relationship between brain and body mass that applies to the moderately sized modern bird species originated in pre-K-Pg times still captures the scaling relationship that applied to relatively small as well as very large Mesozoic theropods, in an extreme example of how biological scaling can be very well described across several orders of magnitude by power functions, which indicate the presence of scale-free, modular, self-organizing principles at work (Herculano-Houzel et al., 2016; West, 2017). Importantly, the overlap between extinct theropods and extant pre-K-Pg birds strongly suggests that theropods were endothermic. Similarly, I find that the brain x body mass scaling relationship of most species of sauropodmorph and ornithischian dinosaurs, as well as pterosaurs, is captured by the scaling relationship that applies to modern ectothermic sauropsids, which suggests that those species of dinosaurs were ectothermic, or if they were endothermic, they had not yet expressed the enlarged brain mass that is presumably affordable by the increased oxidative capacity that powers endothermy (Herculano-Houzel, 2022). However, it remains possible that the values presented here for sauropodmorphs and ornithischians are underestimated, since they stem from a dataset that considered that the brain of these species only filled 50% of the cranial cavity (Hurlburt, 1996); likewise, it is possible that the values for theropods are slightly overestimated, since they assume that in these species, the brain fills the cranial cavity (Balanoff et al., 2013; Witmer et al., 2003).

On a related note, I acknowledge that the nomenclature and quite possibly some of the data provided in Table 1 from the original data of Hurlburt (1996) are outdated, especially those pertaining to sauropodmorphs and ornithischians. I provide those data as initial estimates for non-Theropod species, and maintain the outdated names as in the original dataset, so that those data can be easily located by future researchers, whom I hope will carry this study forward with modern data.

Another methodological issue that will be disputed by some readers is that the scaling relationships reported here are not “corrected” for phylogenetic relatedness, as detailed in the Methods. The central issue of the present study, as in all my previous work, is the scaling relationship between physical quantities such as brain structure mass and numbers of cells and how one predicts the other in different ways in different clades. Whereas phylogenetic comparative methods were initially introduced to alleviate issues with bias possibly introduced by evolutionary relatedness in a sample, I have previously shown that any differences in the scaling exponents calculated with and without accounting for phylogenetic relatedness differ by only 1-2%, which is not a statistically significant difference, provided that there is no mixing in the sample of species from clades already found to scale differently (Gabi et al., 2010; Herculano-Houzel et al., 2011). Moreover, phylogenetic relationships are constantly being updated as research progresses, which would affect the predictive scaling rules calculated if relatedness within each clade were taken into consideration. This is another reason why I continue to prefer to report only the scaling relationships that apply to the raw data (Herculano-Houzel et al., 2015a). These mathematical functions will be readily useful to researchers interested in the relationships between the physical entities involved; other researchers interested in the phylogenetic signal possibly contained within the raw data provided are welcome to apply in their own studies to further phylogenetic studies of evolution, which are not at all the focus of the present work.

A still growing number of studies have focused on establishing whether dinosaurs were ectotherms, mesotherms, or had the high metabolic rates characteristic of modern warm-blooded animals, and the fast-paced behavior and life history that come with it (Werner and Griebeler, 2014; Grady et al., 2014; Wiemann et al., 2022). The present findings on the diverse scaling of brain x body mass across dinosaur clades, which are compatible with endothermy in some and ectothermy in other species, add to the still-going debate about the metabolic condition of fossil dinosaurs by disputing the claim of homogeneous mesothermy across species (Grady et al., 2014) in favor of much larger diversity than previously suspected, supporting the finding that higher metabolic rates appeared in some but not all dinosaur clades (Wiemann et al., 2022). Specifically, while most theropods and the single sauropodmorph (*Diplodocus*) tested had advanced lipoxidation end-products accumulated in quantities indicative of high metabolic rates, different ornithischian and pterosaur species showed concentrations compatible with either high or low metabolic rates (Wiemann et al., 2022). So much diversity amongst dinosaurs and pterosaurs in both metabolism and brain x body scaling (Figure 2b) warrants discontinuation of the practice of treating these species as a mixed bag in scaling studies. Instead of using all-encompassing scaling rules such as the power function shown in Figure 2c, clade-specific analyses and scaling rules should be employed, informed by other features such as analysis of metabolites (Wiemann et al., 2022), which suggests the clustering indicated by the colors in Figure 2c, or by the scaling relationship between brain and body mass, which suggests the clustering indicated by the colors in Figure 2d.

As modeling techniques based on micro-CT data improve and allow the volume of other brain structures to be estimated (Knoll et al., 2021), more evidence should help distinguish which dinosaur and pterosaur species were ecto- or endothermic. The cerebellum, for example, is decidedly larger in extant endothermic species compared to ectothermic species of similar body mass (Kverkova et al., 2022); thus, the size of the cerebellum relative to the mass of the telencephalon (Herculano-Houzel, 2022) and of the body may serve as a new diagnostic criterion to infer the metabolic status of species of the prehistoric fauna. Importantly, this is not a difference of 10-20%, but of 10-20 times in volume of the cerebellum relative to the brainstem between living endothermic and ectothermic amniotes; therefore, models and simulations that estimate the volume of the cerebellum in fossil animals of unknown metabolic status would greatly contribute not only to understanding cerebellar evolution, but also to determining their metabolic status. Still, absent volumetric analyses of the cerebellum, simply determining whether the brain vs body mass relationship clusters with endothermic, pre-K-Pg birds or with ectothermic sauropsids, as more fossil species have their brain and body masses estimated, should already provide diagnostic evidence of the metabolic status of those species.

Estimating the numbers of neurons in the telencephalon, whose main component is the pallium, a major contributor to behavioral flexibility, is obviously consequential for inferring the cognitive capabilities of dinosaur species, whatever their body size (Herculano-Houzel, 2017). The present estimates showing that apex predators such as *Tyrannosaurus* had the numbers of telencephalic neurons found in modern medium-sized primates of impressive cognitive abilities adds a new dimension to how dinosaurs are pictured; an elephant-sized but agile carnivoran biped endowed with macaque- or baboon-like cognition must have been an extremely competent predator indeed. But additionally, I showed recently that the number of neurons in the pallium is a true and reliable predictor of age at sexual maturity and maximal longevity in warm-blooded animals (Herculano-Houzel, 2019a), such that 74% of variation in these life-history variables can be predicted in mammals and birds alike simply by the absolute number of neurons in the cerebral cortex, whereas body mass is an irrelevant predictor once numbers of cortical neurons are accounted for (Herculano-Houzel, 2019a). Using the reported equations L = e^-4.939^ N_cx_^0.402^ and S = e^-2.858^ N_cx_^0.471^ that relate maximal longevity (L) and age at female sexual maturity (S), respectively, to numbers of cortical neurons (Ncx; Herculano-Houzel, 2019), and assuming that most telencephalic neurons in sauropsids are pallial (Kverkova et al., 2022), I can predict that a warm-blooded *Tyrannosaurus* of 2.2-3.3 billion telencephalic neurons would take 4-5 years to reach sexual maturity, and have an estimated maximal longevity of 42-49 years, similar to baboons, whereas *Archaeopteryx* should reach sexual maturity in ca. 8 months, and have a maximal lifespan of 10 years, similar to flycatchers and other songbirds (Herculano-Houzel, 2019). In support of this estimate based on extant warm-blooded species, the survivorship pattern of tyrannosaurs is similar to that seen in long-lived, mammals and birds (Erickson et al., 2006). The predicted sexual maturity of *Tyrannosaurus* at age 5 years, like in modern warm-blooded amniotes of similar numbers of cortical neurons, anticipates by a full decade the previous demonstration that, at 18 years of age, this species was sexually mature (although that was admittedly an upper bound; Lee and Werning, 2008). While the largest and oldest known *T. rex* lived an estimated 28 years, well under the predicted maximal longevity, the finding that only 2% of the population lived long enough to attain maximal size and age for the species (Erickson et al., 2006) makes the estimate of a maximal lifespan of just over 40 years compatible with the oldest known fossil.

Through their association with delayed sexual maturity and longer lifespans, larger numbers of telencephalic neurons simultaneously endow brains with the cognitive flexibility that can be construed as iN_tel_ligence (Herculano-Houzel, 2017) and come with increased lifetime opportunities to develop that increased biological signal processing capability into abilities such as using and creating tools, and devising and perpetuating problem-solving processes (Herculano-Houzel, 2019b). With enough pallial neurons and a long enough lifetime that comes with it, generations overlap enough that developed abilities can be transmitted and perpetuated, forming a body of technology and culture that characterizes populations (Herculano-Houzel, 2019b). The present findings invite the speculation that theropod dinosaurs such as *T. rex*, with even more telencephalic neurons than modern tool-using and tool-making corvids (Olkowicz et al., 2016), had the biological capability to use and craft tools, and develop a culture, like modern birds and primates (Beck, 1974; Whiten et al., 1999; Sapolsky and Share, 2004; von Bayern et al.., 2018). Being able to infer what existed inside the brains of dinosaurs thus multiplies in several directions our knowledge of what life was like in the pre-asteroid, Mesozoic world, and places at least theropods, if not other dinosaurs as well, in the cognitive realm of tool-using and culture-building modern birds and primates.

## Acknowledgements

Thanks to Pavel Nemec and collaborators who collected the reptilian data that made this study possible, and to Fabien Knoll and another reviewer who remained anonymous for greatly improving this manuscript. This work was not supported by any type of funding.

## Competing interests

None.

**All correspondence and material requests should be addressed to the author at the address above.**

## Notes

### Competing Interest Statement

The authors have declared no competing interest.

### Summary of Updates

Revised after peer review

## References

Alvarez LW, Alvarez W, Asaro F, Michel HV (1980) Extraterrestrial cause for the Cretaceous-Tertiary extinction. Science 208, 1095–1108.

Balanoff AM, Bever GS, Rowe TS, Norell MA (2013) Evolutionary origins of the avian brain. Nature 501, 93–97.

Beck BB (1974) Baboons, chimapnzees, and tools. J Human Evol 3, 509–516.

Bennett AF, Ruben JA (1979) Endothermy and activity in vertebrates. Science 206, 649–655.

Bertrand OC, Shelley SL, Williamson TE, Wible JR, Chester SGB, Flynn JJ, Holbrook LT, Lyson TR, Meng J, Miller IM, Püschel HP, Smith T, Spaulding M, Tseng ZJ, Brusatte SL (2022) Brawn before brains in placental mammals after the end-Cretaceous extinction. Science 376, 80–85.

Bininda-Emonds ORP, Cardillo M, Jonew KE, MacPhee RDE, Beck RMD, Grenyer R et al. (2007) The delayed rise of present-day mammals. Nature 446, 507–512.

Boot N, Baas M, van Gaal S, Cools R, de Dreu CKW (2017) Creative cognition and dopaminergic modulation of fronto-striatal networks: Integrative review and research agenda. Neurosci Biobehav Rev 78, 13–23.

Brusatte SL, O’Connor JK, Jarvis ED (2015) The origin and diversification of birds. Curr Biol 25, R888–898.

Erickson GM, Currie PJ, Inouye BD, Winn AA (2006) Tyrannosaur life tables: an example of non-avian dinosaur population biology. Science 313, 213–218.

Gabi M, Collins CE, Wong P, Kaas JH, Herculano-Houzel S (2010) Cellular scaling rules for the brain of an extended number of primate brains. Brain Behav Evol 76, 32–44.

Gillooly JF, McCoy MW (2014) Brain size varies with temperature in vertebrates. PeerJ 2, e301.

Grady JM, Enquist BJ, Dettweiler-Robinson E, Wright NA, Smith FA (2014) Evidence for mesothermy in dinosaurs. Science 344, 1268–1273.

Herculano-Houzel S (2016) What modern mammals teach about the cellular composition of early brains and mechanisms of brain evolution. In Kaas JH, Krubitzer L eds., Evolution of Nervous Systems, 2nd ed., vol. 2, 153–190.

Herculano-Houzel S (2017) Numbers of neurons as biological correlates of cognitive capability. Curr Opin Behav Sci 16, 1–7.

Herculano-Houzel S (2019a) Longevity and sexual maturity vary across species with number of cortical neurons, and humans are no exception. J Comp Neurol 527, 1689–1705.

Herculano-Houzel S (2019b) Life history changes accompany increased numbers of cortical neurons: a new framework for understanding brain evolution. Prog Brain Res 250, 179–216.

Herculano-Houzel S (2022) Mammals, birds and non-avian reptiles have signature proportions of numbers of neurons across their brain structures: Numbers of neurons increased differently with endothermy in birds and mammals. bioRxiv.org, 2022/496835.

Herculano-Houzel S, Lent R (2005) Isotropic fractionator: a simple, rapid method for the quantification of total cell and neuron numbers in the brain. J Neurosci 25, 2518–2521.

Herculano-Houzel S, Kaas JH (2011) Gorilla and orangutan brains conform to the primate scaling rules: implications for hominin evolution. Brain Behav Evol 77, 33–44.

Herculano-Houzel S, Ribeiro P, Campos L, da Silva AV, Torres LB, Catania KC, Kaas JH (2011) Updated neuronal scaling rules for the brains of glires (rodents/lagomorphs). Brain Behav Evol 78, 302–314.

Herculano-Houzel S, Ribeiro PFM, Campos L, da Silva AV, Torres LB, Catania KC, Kaas JH (2011) Updated neuronal scaling rules for the brains of Glires (rodents/lagomorphs). Brain Behav Evol 78, 302–314.

Herculano-Houzel S, Manger PR, Kaas JH (2014) Brain scaling in mammalian brain evolution as a consequence of concerted and mosaic changes in numbers of neurons and average neuronal cell size. Front Neuroanat 8, 77.

Herculano-Houzel S, Catania K, Manger PR, Kaas JH (2015a) Mammalian brains are made of these: a dataset on the numbers and densities of neuronal and non-neuronal cells in the brain of glires, primates, scandentia, eulipotyphlans, afrotherians and artiodactyls, and their relationship with body mass. Brain Behav Evol 86, 145–163.

Herculano-Houzel S, Kaas JH, Miller D, Von Bartheld CS (2015b) How to count cells: the advantages and disadvantages of the isotropic fractionator compared with stereology. Cell Tissue Res 360, 29–42.

Hopson JA (1977) Relative brain size and behavior in archosaurian reptiles. Annu Rev Ecol Systematics 8, 429–448.

Hulburt GR (1996) Relative brain size in recent and fossil amniotes: Determination and interpretation. PhD Thesis, University of Toronto.

Hulburt GR, Ridgely RC, Witmer LM (2013) Relative size of brain and cerebrum in Tyrannosaurid dinosaurs: an analysis using brain-endocast quantitative relationships in extant alligators. In Parrish JM, Molnar RE, Currie PJ, Koppelhus EB (eds), Tyrannosaurid paleobiology. Indiana University Press, Bloomington, 134–154.

Jerison HJ (1973) Evolution of the brain and intelligence. New York, Academic Press.

Knoll F, Buffetaut E, Bülow M (1999) A theropod braincase from the Jurassic of the Vaches Noires cliffs (Normandy, France): osteology and palaeoneurology. Bull Soc Geol France 170, 103–109.

Knoll F, Schwarz-Wings D (2009) Palaeoneuroanatomy of Brachiosaurus. Annales Paléontol 95, 165–175.

Knoll, F., S. Lautenschlager, S. Kawabe, G. Martínez, E. Espílez, L. Mampel, & L. Alcalá. 2021. Palaeoneurology of the Early Cretaceous iguanodont *Proa valdearinnoensis* and its bearing on the parallel developments of cognitive abilities in theropod and ornithopod dinosaurs. Journal of Comparative Neurology, 529 (18): 3922–3945.

Knoll, F., S. Kawabe & A. Watanabe (2022) A proxy for brain-to-endocranial cavity index in non-neornithean dinosaurs and other extinct archosaurs. 10th European Conference on Comparative Neurobiology, Abstract Book. 33. Eds: P. Němec, K. Kverková, Y. Zhang, F. Dionigi, R. Druga & G. Pavlinkova. Praha: Univerzita Karlova.

Ksepka DT, Balanoff AM, Smith NA, Bever GS, Bhullar B-AS, Bourdon E et al. (2020) Tempo and pattern of avian brain size evolution. Curr Biol 30, 2026–2036.

Kverkova K, Marhounová L, Polonyiová A, Kocourek M, Zhang Y, Olkowicz S, Stratková B, Pavelková Z, Vodicka R, Frynta D, Nemec P (2022) The evolution of brain neuron numbers in amniotes. Proc Natl Acad Sci USA 119, e2121624119.

Lee AH, Werning S (2008) Sexual maturity in growing dinosaurs does not fit reptilian growth models. Proc Natl Acad Sci USA 105, 582–587.

Olkowicz S, Kocourek M, Lucan RK, Portes M, Fitch WT, Herculano-Houzel S, Nemec P (2016) Birds have primate-like numbers of neurons in the telencephalon. Proc Natl Acad Sci USA 113, 7255–7260.

Rowe TB, Macrini TR, Luo Z-X (2011) Fossil evidence on origin of the mammalian brain. Science 332, 955–957.

Sapolsky RM, Share LJ (2004) A pacific culture among wild baboons: its emergence and transmission. PLoS Biol 2, e106.

Sol D, Olkowicz S, Sayol F, Kocourek M, Zhang Y, Marhounová L, Osadnik C, Corssmit E, Garcia-Porta J, Martin TE, Lefebvre L, Nemec P (2022) Neuron numbers link innovativeness with both absolute and relative brain size in birds. Nature Ecol Evol, doi.org/10.1038/s41559-022-01815-x.

Ströckens F, Neves K, Kirchem S, Schwab C, Herculano-Houzel S, Güntürkün O (2022) High associative neuron numbers could drive cognitive performance in corvid species. J Comp Neurol 530, 1588–1605.

Von Bayern AMP, Danel S, Auersperg AMI, Mioduszewska B, Kacelnik A (2018) Compound tool construction by New Caledonian crows. Scientific Reports 8, 15676.

Werner J, Griebeler EM (2014) Allometries of maximum growth rate versus body mass at maximum growth indicate that non-avian dinosaurs had growth rates typical of fast growing ectothermic sauropsids. PLoS One 9, e88834.

West G (2017) Scale: The universal laws of growth, innovation, sustainability, and the pace of life in organisms, cities, economies, and companies. Penguin Press, New York.

Whiten A, Goodall J, McGrew WC, Nishida T, Reynolds V, Sugiyama Y, Tutin CEG, Wrangham RW, Boesch C (1999) Cultures in chimpanzees. Nature 399, 682–685.

Wiemann J, Menéndez I, Crawford JM, Fabbri M, Gauthier JA, Hull PM, Norell MA, Briggs DEG (2022) Fossil biomolecules reveal an avian metabolism in the ancestral dinosaur. Nature, https://doi.org/10.1038/s41586-022-04770-6.

Williams RW, Herrup K (1980) The control of neuron number. Annu Rev Neurosci 11, 423–253.

Witmer LM, Chatterjee S, Franzosa J, Rowe T (2003) Neuroanatomy of flying reptiles and implications for flight, posture and behaviour. Nature 425, 950–953.

Yu Y, Zhang C, Xu X (2021) Deep time diversity and the early radiations of birds. Proc Natl Acad Sci USA 118, e2019865118.

